# Improving genomic selection in hexaploid wheat with sub-genome additive and epistatic models

**DOI:** 10.1101/2024.04.19.590350

**Authors:** Augusto Tessele, David O. González-Diéguez, Jose Crossa, Blaine E. Johnson, Geoffrey P. Morris, Allan K. Fritz

## Abstract

The goal of wheat breeding is the development of superior cultivars tailored to specific environments, and the identification of promising crosses is crucial for the success of breeding programs. Although genomic estimated breeding values were developed to estimate additive effects of genotypes before testing as parents, application has focused on predicting performance of candidate lines, ignoring non-additive genetic effects. However, non-additive genetic effects are hypothesized to be especially importance in allopolyploid species due to the interaction between homeologous genes. The objectives of this study were to model additive and additive-by-additive epistatic effects to better delineate the genetic architecture of grain yield in wheat and to the improve accuracy of genomewide predictions. The dataset utilized consisted of 3740 F_5:6_ experimental lines tested in the K-State wheat breeding program across the years 2016 and 2018. Covariance matrices were calculated based on whole and sub-genome marker data and the natural and orthogonal interaction approach (NOIA) was used to estimate variance components for additive and additive-by-additive epistatic effects. Incorporating epistatic effects in additive models resulted in non-orthogonal partitioning of genetic effects but increased total genetic variance and reduced deviance information criteria. Estimation of sub-genome effects indicated that genotypes with the greatest whole genome effects often combine sub-genomes with intermediate to high effects, suggesting potential for crossing parental lines which have complementary sub-genome effects. Modeling epistasis in either whole-genome or sub-genome models led to a marginal (3%) but significant improvement in genomic prediction accuracy, which could result in significant genetic gains across multiple cycles of breeding.

## Introduction

In wheat breeding, selection of experimental lines has historically been based on phenotypic selection, which encapsulates all forms of genetic effects. However, utilization of genomic selection (Meuwissen et al. 2001) to predict the performance of candidate lines overlooks the potential contribution of non-additive effects, especially epistasis, which potentially could play an important role in the genetic expression of agronomic traits (Chapman and McNeal, 1971; Sun et al., 1972; Goldringer et al., 1997; Zhang et al., 2008; Jiang et al., 2017). Epistasis is defined as the interaction between alleles of different genes and is normally classified into additive-by-additive, additive-by-dominance, and dominance-be-dominance, with higher order interactions normally ignored.

Historically, complex mating designs were used to create the progeny that could be used to estimate epistatic effects, e.g., the triple test cross (Bauman, 1959). Current breeding programs have an abundance of experimental lines genotyped with phenotypic records that could be leveraged to estimate genetic effects. With large number of markers covering the genome and quantitative trait loci assumed to have a normal distribution, genome-wide markers can be used to estimate the additive covariance between individuals and predict their additive genetic value (Nejati-Javaremi et al. 1997; VanRaden 2008). To obtain an approximation of the additive-by-additive relationship matrix between individuals, Henderson (1985) proposed the use of the Hadamard product of the additive covariance matrix based on pedigree information, which was later expanded to accommodate genomic-based additive relationship matrices (Su et al., 2012; Jiang and Reif, 2015). Newly, the natural and orthogonal interaction (NOIA) approach (Alvarez-Castro and Carlborg, 2007) was expanded to build genomic relationship matrices using genotypic frequencies and to include genome-wide epistasis (Vitezica et al., 2017), resulting in an orthogonal partitioning of genetic variance components, even if not in Hardy-Weinberg Equilibrium (HWE). The main feature of the NOIA approach is to account for Hardy-Weinberg disequilibrium, which is quite common in agriculture or livestock e.g. inbred line populations, F_1_ crosses, three-way crosses, or backcrosses. However, most of the previous studies on estimation of genomic variance have ignored this by assuming HWE. If genomic relationship matrices are built incorrectly assuming HWE, inclusion of more genetic effects in the model can dramatically change estimates (Vitezica et al., 2017). In the absence of HWE, and to get meaningful estimates of variances that sum to the phenotypic variance, special care in standardization of covariance matrices may be needed (Legarra 2016), and this has not been considered by other authors. Standardization consists of scale the covariance matrix to have average diagonal equal to 1, and mean of the matrix equal to zero (Legarra 2016). Moreover, to address the challenge of partitioning estimates of genomic variance in the presence of extensive linkage disequilibrium (LD), as is the case of common wheat, Sorensen, Fernando, and Gianola (2001) and Lehermeier et al., 2017, developed methods to account for the contribution of the covariance between loci, i.e., linkage disequilibrium and covariance between genetic effects, when estimating genomic variances. This approach allows variance and covariance to contribute to genetic effects, thereby preventing the overestimation or underestimation of these effects. Thus, there is the potential to better delineate the genetic architecture of important agronomic traits while leveraging existing data sets in crop species, as demonstrated on wheat grain yield using historical CIMMYT and Cornell wheat breeding data (Santantonio et al., 2019) and on important wheat diseases using synthetic wheat data (Cuevas et al., 2024).

Common wheat (*Triticum aestivum* spp *aestivum*) is a hexaploid species originated from the interspecific hybridization between tetraploid wheat, *Triticum turgidum* (Sax 1922; Kihara 1924), the source of the AABB genome, and *Aegilops tauschii* (Kihara 1944; McFadden and Sears, 1946), the source of the DD genome. The presence of homeologous genomes could create positive or negative epistasis through subfunctionalization (Lynch and Force, 2000) or substrate competition (Qian et al., 2010), while also impacting gene expression patterns (Leach et al., 2014; Akhunova et al., 2010). Intra-genomic interactions have also been reported in wheat (Tranquilli and Dubcovsky, 2000; Reif et al., 2011; Sehgal et al., 2020), underscoring the potential importance of epistasis underpinning agronomic traits. In addition, the inbreeding nature of wheat leads to a fast fixation of alleles (Charlesworth, 2003), revealing epistatic interactions whose favorable combinations are perpetuated with selection of superior genotypes in breeding programs. Although hexaploid, the three sub-genome of common wheat undergo disomic segregation (Feldman and Levy, 2012) and are normally treated as diploids in breeding programs. For that reason, partitioning whole genome genetic effects into sub-genome opens up the possibility to attribute biological importance to whole genome effects and variance estimates, as was previously done by Santantonio et al., (2019), Cuevas et al., 2024 and Bernardo (2020).

One of the challenges in accurately estimating the extent of epistatic effects in wheat lines in the conversion of epistatic variance into additive variance, which reaches its maximum when an allele becomes fixed (Whitlock et al., 1995; Cheverud and Routman, 1996; Holland, 2001; Technow et al., 2019). The Kansas State wheat breeding program is notable for maintaining a large genetic base through consistent crossings with external germplasm and wild accessions. This breeding strategy enhances the probability of observing epistatic interactions segregating while increasing the minor allele frequency of such interactions. Consequently, the statistical power to detect epistatic effects is higher compared to studies that relied in less diverse breeding data (Santantonio et al., 2019). Therefore, the objectives of this study were to: (i) partition the genetic variance of wheat grain yield into additive and epistatic effects; (ii) partition whole genome effects into sub-genome effects; and (iii) incorporate epistasis into the genomic selection model.

We used a population of hexaploid wheat genotypes from the Kansas State wheat breeding program and implemented recent developments and methods for estimating genetic variance components as the NOIA approach and correction for LD and covariance between genetic effects.

## Material and Methods

### Estimation of Epistatic Variance

#### Phenotypic Data

Phenotypic data consisted of grain yield records for 3740 F_5:6_ wheat genotypes grown in the early yield trail stage, comprised of individual plan short rows (IPSR), of the K-State hard red winter wheat breeding program across the years of 2016 and 2018. The IPSR is the first stage of the breeding program where selection is based on grain yield data (previous stages consisted of visual single plant selection). The IPRS represents the foundational genetic diversity within each breeding cycle. The experimental design used for the phenotypic evaluation consisted of unreplicated trials and was a modified augmented design (Federer and Raghavarao, 1975) type II with one replication of each experimental line. However, to quantify and remove spatial variability of these large experiments, we employed a two-stage analyses (Smith et al., 2001; Welham et al., 2010), and included fitting a spatial model to account for field patterns associated with environmental factors (Cullis and Gleeson, 1991; Cullis et al., 1998; Smith et al., 2001). Due to the limited number of replications in the trials, an extra step was added to the first-stage analysis, as reported in (Arief et al., 2019). In the first step, the genotypes were fitted as random effects to provide an estimate of the spatial trend, which permits utilizing all genotypes to estimate the trend. In the second step, the genotypes were fitted as fixed effects to calculate the best linear unbiased estimates (BLUEs) necessary for the second-stage analysis (Smith et al., 2001). The statistical model used in both steps of the first-stage analysis was:

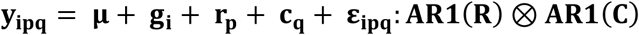

Where **y**_**ipq**_ is the grain yield of genotype i in row p and column q; **μ** is the intercept; **g**_**i**_ is the effect of genotype; **r**_**p**_ is the random effect of row p; **c**_**q**_ is the random effect of column q; and **ε**_**ipq**_ is the residual effect modeled using first-order autoregression (AR1) for row q and column q. As previously described, the initial phase of our analysis employed the first-step approach to calculate variance component estimates for the random spatial terms, while fitting the genotype as a random effect. In the second step, these variance component estimates were utilized to derive the BLUEs for the genotypes and their corresponding weights, where genotype was treated as a fixed effect. The weights were computed as the inverse of the residual variance for each field, multiplied by the replication count for each genotype, and the pooled residual variance across all fields (Cullis et al. 1996).

The second-stage analysis was conducted using the BLUEs and weights for genotypes in order to obtain the BLUPs according to the following statistical model:

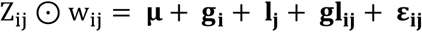

Where Z_ij_ ⊙ w_ij_ is the weighted BLUE from the first-stage analysis for genotype i in location j; **l**_**j**_ is the effect of location j; **gl**_**ij**_ is the effect of genotype i in location j; and **ε**_**ij**_ is the residual. The BLUPs obtained from the second-stage analysis were used in the downstream analyses of this study. The two-stage analysis was conducted using the Echidna Mixed Model Software (Gilmour, 2018).

#### Genotypic Data

The lines analyzed were in the F_5:6_ generation and were genotyped using genotype-by-sequencing (GBS) (Poland et al. 2012). To ensure data quality, markers with a minor allele frequency lower than 0.01, more than 20% missing values, and more than 20% heterozygotes were filtered out. The marker imputation process was carried out using the ‘rrBLUP’ package (Endelman, 2011), where missing markers were imputed as the mean value among all lines for that marker, resulting in a dataset containing a total of 55,148 single nucleotide polymorphisms (21,338 SNPs on sub-genome A, 26,996 on sub-genome B, and 6,844 for sub-genome D).

### Statistical Models

Four models were used to quantify additive and additive-by-additive epistatic (herein described as epistasis) effects across sub-genomes. The first two models consisted of analyses at the whole genome level while the last two models partitioned genetic effects, both additive and epistatic, by sub-genome. Variance components, for all models, were estimated using the natural and orthogonal interaction approach (NOIA) parametrizations outlined by Alvarez-Castro and Carlborg (2007) and Vitezica et al. (2017).

### Whole Genome Models

#### Model 1 – Additive Effects

The linear model for grain yield that includes only an additive term is given by:

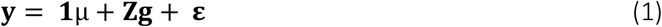

where **y** is the response vector (adjusted phenotypic data), **μ** is the intercept, **ε** is a vector of random residuals with 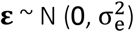, where 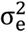is the residual variance, **Z** is a design matrix of random effects and **g** is the vector of additive genomic breeding values with 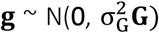 where 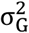 is the genomic additive genetic variance and the **G** is an additive genomic relationship constructed according to Vitezica et al. (2017) (see Raffo et al. 2022).

#### Model 2–-Additive and Epistatic Effects

Following Henderson (1985), the epistatic covariance of individuals can be calculated as the Hadamard product of the component covariance matrices. Hence, the second model used in this study extends the first model by including an additive-by-additive epistatic term:

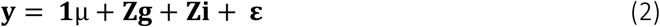

where **y**, **μ**, **Z**, **g** and **ε** are the same as in the first model, **i** is a vector of additive-by-additive epistatic genomic effects with 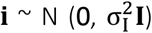, where 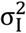 is the epistatic variance and **I** is the additive-by-additive relationship matrix, built following Vitezica et al. (2017). This method consists of using the Hadamard product of the additive genomic relationship matrix, scaled to have an average diagonal equal to 1, to provide meaningful variance estimates (Legarra 2016) at the whole genome (and subsequently at the sub-genome levels). The matrices are designed to capture deviations from additivity due to epistatic interactions across loci, thereby accounting for non-additive genetic variance arising from epistatic deviation effects.

### Sub-Genome Models

#### Model 3 – Additive Effects

As described in Santantonio et al (2019), **G**_**j**_can be decomposed into individual additive effects for each sub-genome, such that **G**_**j**_ **=A**_**j**_ **+ B**_**j**_ **+ D**_**j**_, resulting in the following model:

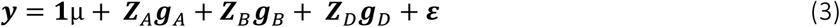

Decomposition of the whole genome into sub-genomes permits each sub-genome to have an individual additive genetic variance and covariance among individuals, such that 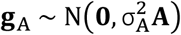, 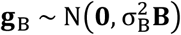 and 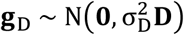. The matrices **A, B**, and **D** are the realized additive genetic variance-covariance for each sub-genome and are calculated using the method used for calculating the whole genome additive variance-covariance matrix.

#### Model 4 – Additive and Epistatic Effects

The whole genome epistatic component, **I**, can also be decomposed into intra and inter subgenome epistatic interactions, such that **I**_**i**_ = **AA**_**i**_ **+ BB**_**i**_ **+ DD**_**i**_ **+ AB**_**i**_ **+ AD**_**i**_ **+ BD**_**i**_. The new additive plus epistatic sub-genome model is then expressed as:

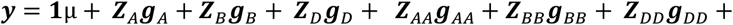

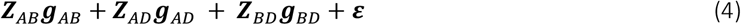

Decomposition of the whole genome epistatic term into sub-genomes results in expression of additive-by-additive genetic variance-covariance among individuals for each pair of sub-genomes. This decomposition is generalized as 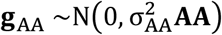, and 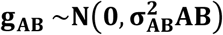, where the matrices **AA**,and **AB**, are the realized intra and inter sub-genome epistatic variance-covariance matrices, respectively, calculated using the previously described procedures.

#### Variance Components, Covariance, and Heritability

Variance components were estimated using the Bayesian BGLR R-package (Pérez and de los Campos, 2014). To enhance computational efficiency, eigenvalue decomposition of the variance-covariance matrices described by Acosta-Pech et al. (2017), was used and modeled as Bayesian Ridge Regression. For each model, a total of 50,000 iterations were executed, with the first 5,000 iterations discarded as burn-in. Subsequent iterations were thinned by selecting one of every ten samples, resulting in 4,500 samples used for estimating the variance components. Based upon a Bayesian Markov Chain Monte Carlo (MCMC) framework, genetic variance components were calculated using the genetic values derived from each MCMC sample, per the methods of Sorensen, Fernando, and Gianola (2001). By following this method, we account for the contribution of the covariance between loci, i.e., linkage disequilibrium when estimating genomic variances (Lehermeier et al., 2017). As described by Lehermeier et al. (2017), within a Bayesian framework, samples from the posterior distribution of the total genetic variance, incorporating covariances among effects, can be derived from posterior samples of the effects themselves. In this study, following ideas presented by Lehermeier et al. (2017), variance components from each MCMC sample were used to calculate the correlation between additive and epistatic terms in both whole genome (Model 2) and sub-genome models (Model 3 and 4). Broad-sense heritability was calculated for each model as the ratio of total genetic variance divided by total phenotypic variance.

#### Genomic prediction

Five-fold cross validation, with ten replicates was used to test predictive ability of the models. For each replicate, the set of 3740 lines was randomly divided into five groups, irrespective of which year they were tested, with four used to train the model and the remaining used to predict genetic values. Using that structure, all four above-described models were fitted, resulting in predicted genetic values for each line. Prediction of the five folds was correlated with the phenotypic value, with the resulting value representing the prediction accuracy of the model. Correlation between predicted genetic values and phenotypic values provided comparisons of accuracy amongst models. To test statistical significance between the prediction accuracies of the models fitted, Tukey’s honestly significant difference was calculated using the ‘agricolae’ package on R (de Mendiburu, 2019).

## Results

### Variance Components and Broad-Sense Heritability

#### Whole Genome Models 1 and 2

Estimated posterior means of the whole genome additive Model 1 resulted in additive genetic effects accounting for 27% of the observed phenotypic variance for wheat grain yield. However, in whole genome additive plus epistatic Model 2, the contribution of additive variance decreased to only 11%, while epistatic variance became prominent, explaining 29% of the observed phenotypic variance. By including an epistatic term in the whole genome additive plus epistatic Model 2, total genetic variance increased while residual variance decreased compared to whole genome additive Model 1 (Figure 1). These results suggest that the epistatic term, although not orthogonal to additive effects, captures a portion of the genetic variance that is not captured by the additive term alone. Consequently, broad-sense heritability estimated using the whole genome additive plus epistatic Model 2 (0.39) exceeded the estimate of the fully additive model (0.27 – see Supplemental Table 1). Interestingly, while epistatic effects did not result in orthogonal partitioning of genetic effects, the correlation between additive and epistatic terms was virtually zero (Supplemental Table 2). The deviance information criteria value was smaller for whole genome additive plus epistatic Model 2 than for whole genome additive Model 1 (Supplemental Table 1), suggesting that the incorporation of an epistatic term captured epistatic interactions and provide a better fit of data.

**Figure 1.**
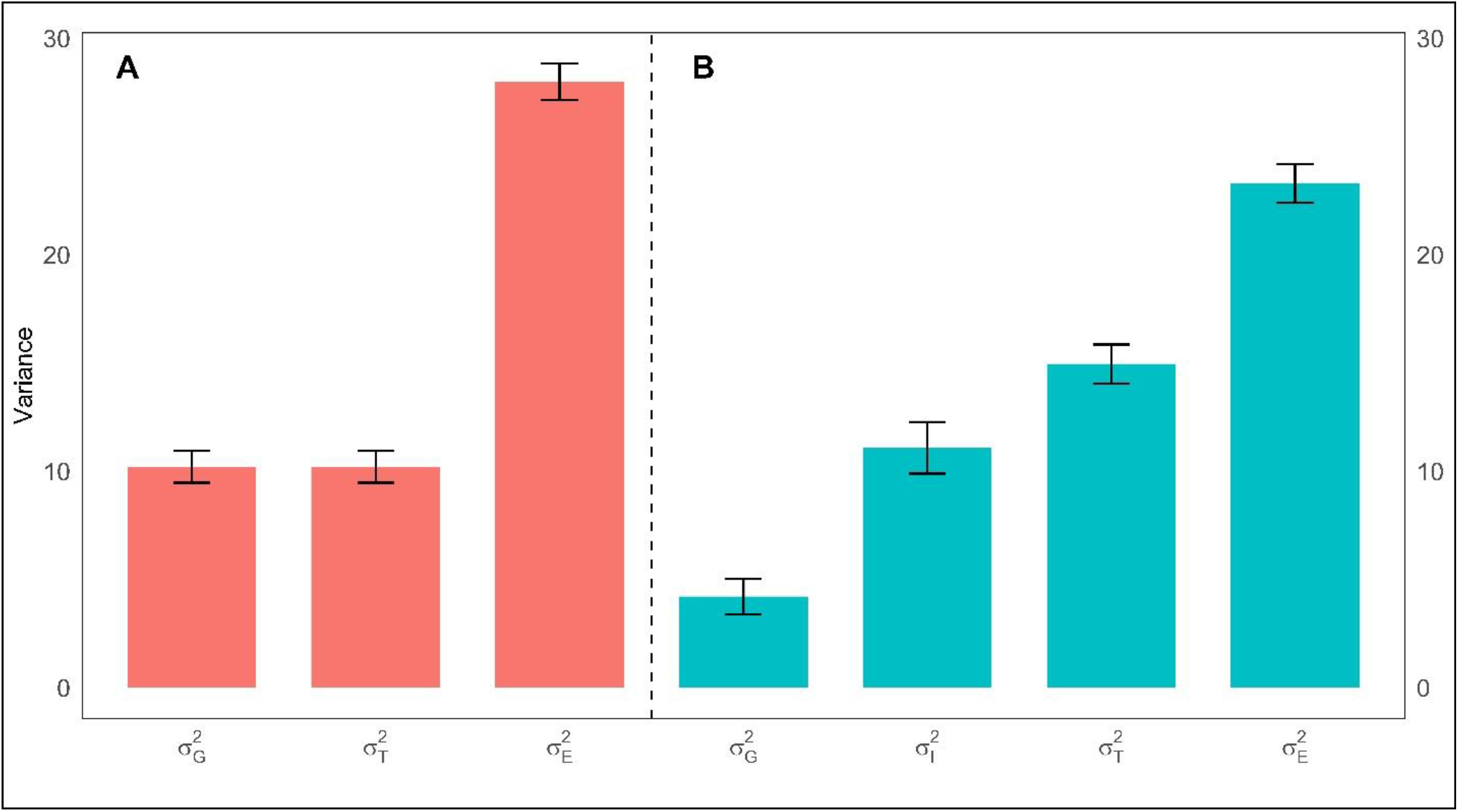
Substantial additive-by-additive epistatic contribution to variance of grain yield. Estimated posterior means and standard deviations of genetic variance components of Model 1 (panel A) and Model 2 (panel B). 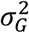 refers to additive variance, 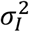 to additive-by-additive epistatic variance, 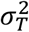 to total genetic variance, and 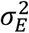 is the residual variance of the model.

#### Sub-Genome Models

The sum of additive genetic variances across genomes for the sub-genome additive Model 3 (10.88) was similar to the total genetic variance estimated using the whole genome additive Model 1, which was expected since breaking the genome into sub-genomes should not imply different additive effects (Figure 1 and 2). For the sub-genome additive plus epistatic Model 4, the sum of sub-genome additive effects added to 4.6 and the epistatic variance was 12.7, indicating a slight increase in epistatic variance with little change in additive variance compared to corresponding values from the whole genome additive plus epistatic Model 2 (Supplemental Table 3). The sum of the additive and epistatic variance deviated from the observed total genotypic variance because of a negative covariance between the terms in the sub-genome additive plus epistatic Model 4 (Supplemental Table 6). Using sub-genome additive Model 3, the largest additive variance was estimated for the A sub-genome, followed by the corresponding estimates for the D and B sub-genomes (Figure 2). Inclusion of the intra- and inter-sub-genome interaction terms reduced the amount of additive variance associated with each sub-genome, and the D sub-genome displayed the highest additive effects among the three sub-genomes. As with whole genome additive plus epistatic Model 2, the addition of intra and inter sub-genome epistatic terms resulted in non-orthogonal partitioning of genetic effects. Likewise, the correlation within and between additive and epistatic terms was nearly zero in all instances (Supplemental Table 6).

**Figure 2.**
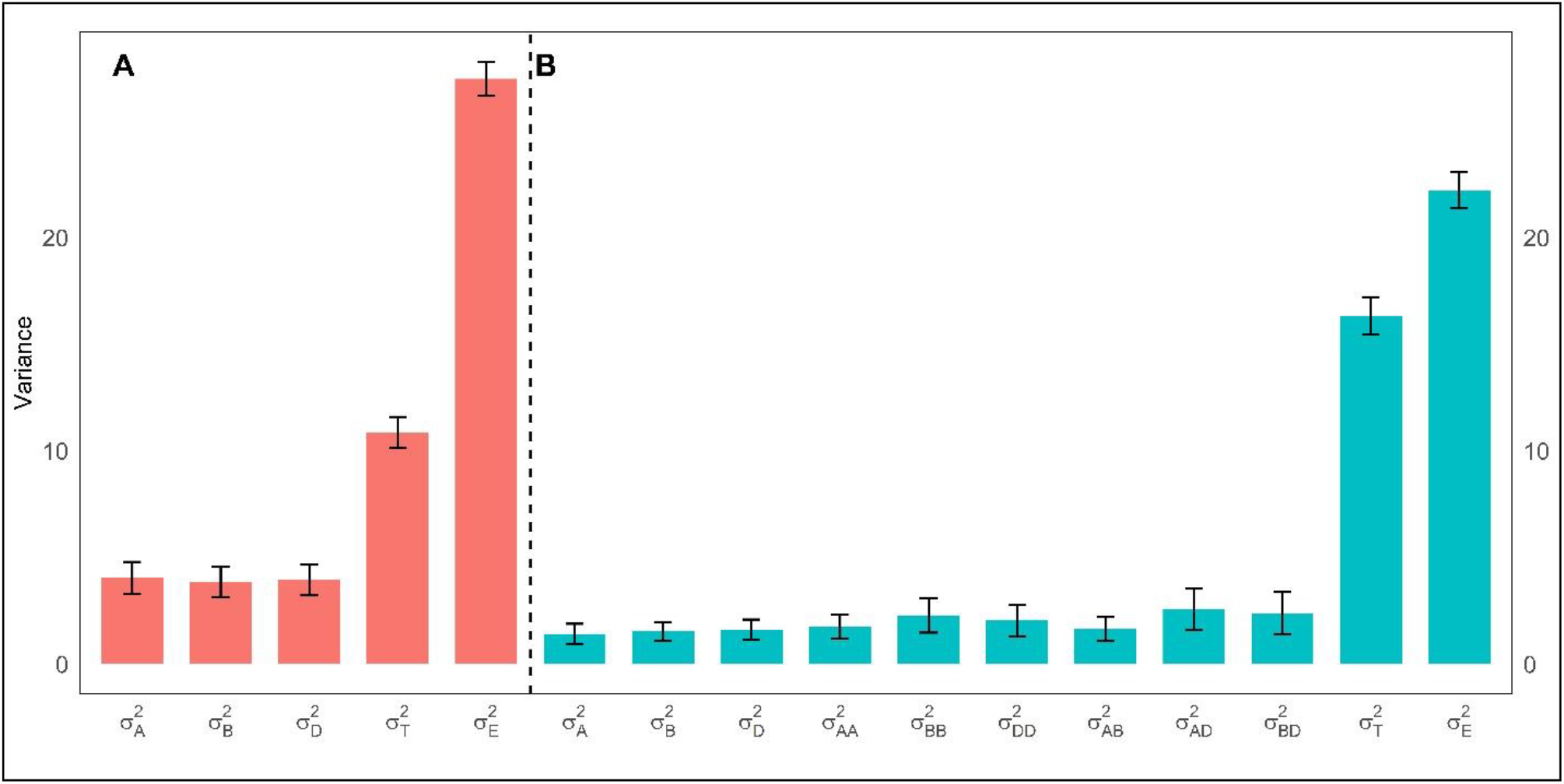
Wheat sub-genomes additive and epistatic variance components in grain yield. Estimated posterior means and standard deviations of genetic variance components of Model 3 (panel A) and Model 4 (panel B). 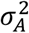 refers to additive variance in subgenome A, 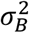 to subgenome B and 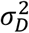 to D. Intra and inter sub-genomes additive-by-additive epistatic variance are depicted by 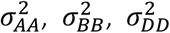, and 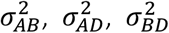, respectively. 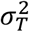 to total genetic variance, and 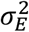 is the residual variance of the model.

The larger total genetic variance estimated using Model 4 increased the magnitude of estimated broad-sense heritability compared to estimates from the whole genome Models 1 and 2 (Supplemental Table 3). In addition, the deviance information criteria values were smaller than those obtained using the whole genome models, indicating a better fit of data.

### Correlation Between Whole and Sub-Genome Breeding Values

Correlations between whole and sub-genomes effects were estimated to characterize the relationships between sub-genome additive effects and whole genome effects (Figure 3). The smallest correlation was observed between the effects of sub-genome D and the whole genome effects (0.61 – Supplemental Table 4), while the correlations between the effects of sub-genome A and B with the whole genome effects were higher (0.72 and 0.69, respectively). The correlations amongst sub genome effects was very low. Interestingly, when analyzing the sub-genome additive effects from Model 4, which includes epistasis, the correlations between sub-genome effects and whole genome effects, as well as between sub-genomes, generally decreased, except for the correlation between sub-genomes A and D. Although correlations between sub-genomes and whole genome varied, all three sub-genomes resulted in similar maximum and minimum additive effects, and standard deviations (Supplemental Table 5).

**Figure 3.**
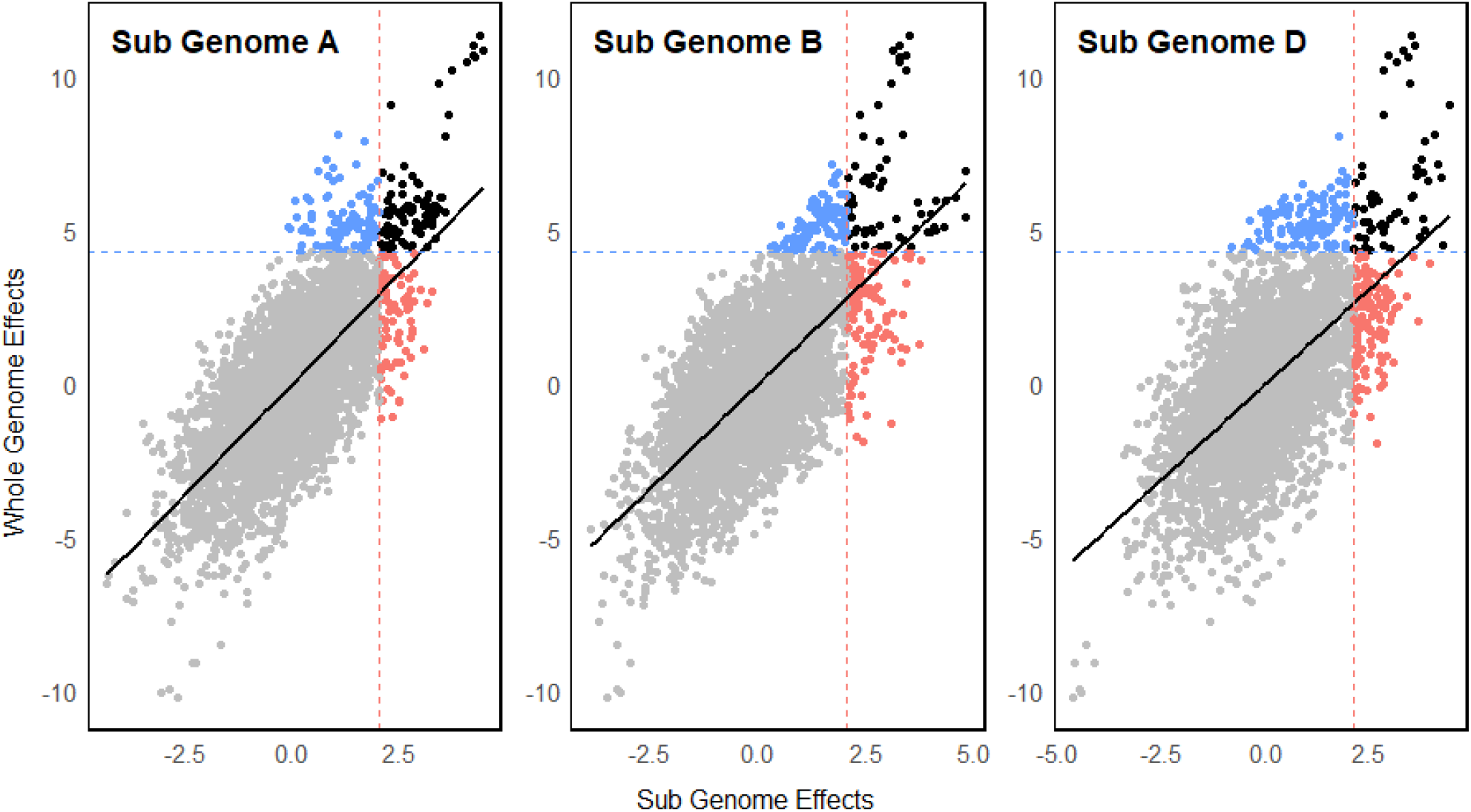
Sub-genome additive effects correlates with whole genome effects for grain yield in wheat. The dotted lines indicate the 95 percentile for whole or subgenome additive effects. The solid black lines indicate the correlation between whole genome and sub genome effects. Blue dots indicate genotypes with whole genome additive effects above the 95-percentile. Pink dots indicate genotypes with sub genome additive effects above the 95-percentile. Black dots indicates genotypes above the 95-percentile for both thresholds.

In total, 98 lines were found above the 95-percentile threshold for both whole and sub-genome A additive effects (Figure 3). Of those, 28 and 22 genotypes were also above the 95-percentile threshold for sub-genome B and D, respectively. In sub-genome B and D, 75 and 57 lines were found above the 95-percentile threshold for both sub-genomes and whole genome additive effects, with 22 lines above the 95-percentile threshold for both sub genomes B and D. In total, 11 genotypes were above the 95-percentile for all sub-genomes, 9 of which were sister lines. Hence, in general, genotypes with the highest whole genome effects tended to also exhibit intermediate to high additive effects in more than one sub-genome. Of the top-100 genotypes with highest whole genome additive effects, 48 were advanced to the preliminary yield trials (PYT). Among these, 6 were selected for the advanced yield trials (AYT), but none of these genotypes were advanced to the elite trials (EYT). One possible reason for the discarding of genotypes with high whole genome additive effects could be selection for or against other important agronomic traits not considered in this prediction model, traits such as disease resistance, maturity, and lodging. For example, from this pool of 3740 genotypes evaluated across three breeding cycles, only eight reached the EYT stage, five of which were above the top-90 percentile. It is necessary to acknowledge that the total genetic value of each genotype across many traits is used to make advancement decisions, with those genetic values being a function of both additive and epistatic effects. In that regard, the sub-genome additive plus epistatic Model 4 estimated higher genotypic values for the eight lines that reached the EYT stage, with seven of those eight falling withing the top-10 percentile. Compared to the sub-genome additive Model 3, these eight lines had an average rank change of 127 positions towards the top when epistasis was modeled. This result of selection highlights the importance of epistasis when making selection decisions based upon the genetic value of candidate genotypes.

### Genomic Predictive Ability

The prediction accuracy between the predicted total genetic values and the phenotypic values was evaluated using the five-fold cross validation method (Figure 4). For both whole and sub-genome models, inclusion of epistatic genetic effects significantly improved the prediction accuracy compared to fully additive models, based upon Tukey’s honestly significant difference test. In contrast, partitioning whole genome effects into sub-genome effects did not improve the accuracy of predictions. The correlations amongst total genetic values of the four models fitted showed very high values (Supplemental Table 7). As expected, the smallest correlation was observed between whole genome additive Model 1 and sub-genome additive plus epistatic Model 4. Similarly, correlations amongst genomic estimated breeding values (sum of additive effects only) included a smaller correlation between the purely additive and the additive plus epistatic models (Supplemental Table 8).

**Figure 4.**
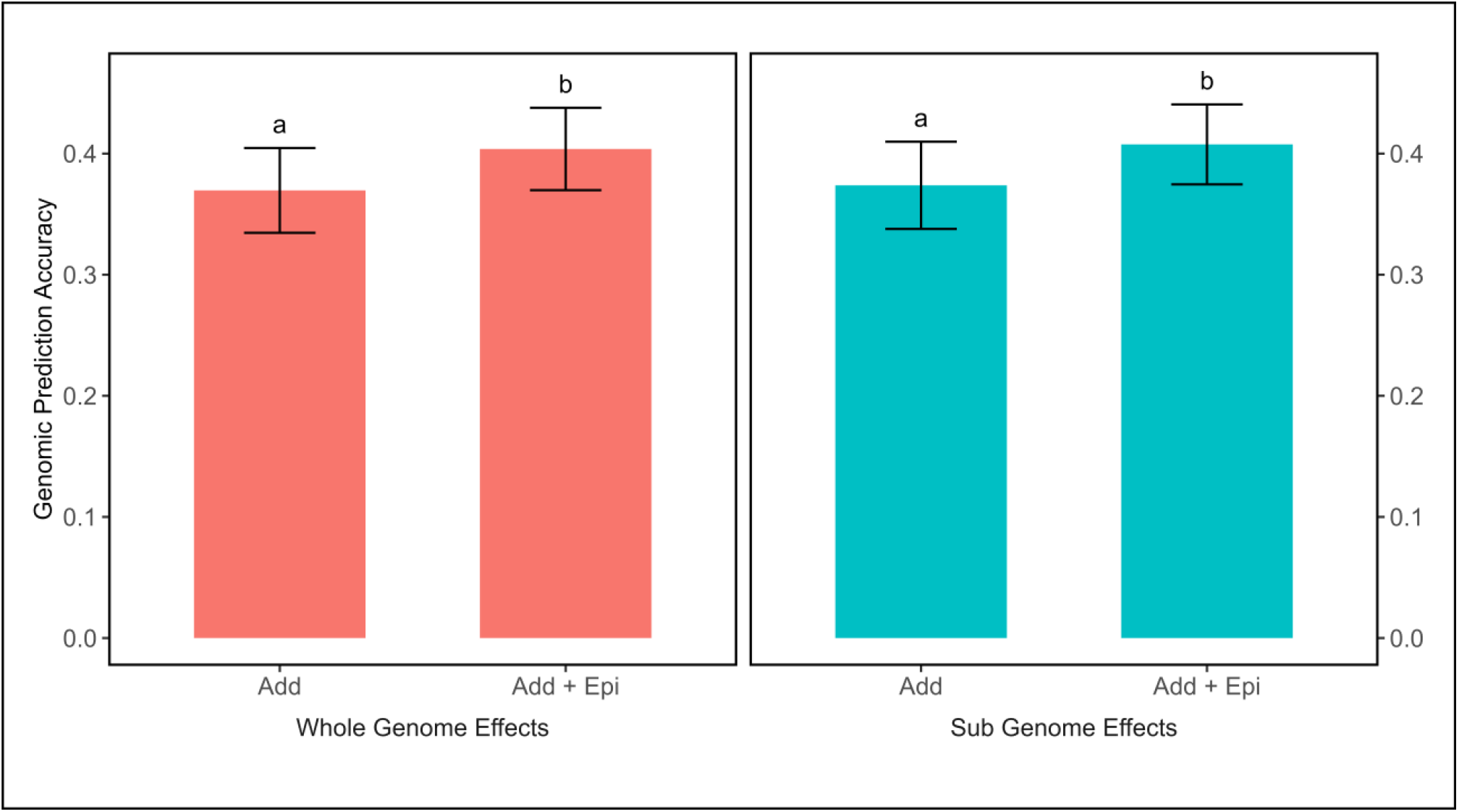
Increased prediction accuracy using models that include epistasis. Bar plot of predictive abilities of additive whole (panel A) and sub-genome (panel B) models with the inclusion or not of an epistatic term using a five-fold cross validation appraoch with 10 replications.

## Discussion

### Modeling Epistasis Improves the Total Genetic Variance and Broad-Sense Heritability

The role of epistasis is expected to be especially relevant in polyploid species due to the interaction between homologous genes, but epistasis has been commonly ignored in wheat. In this study, based upon modeling variance components, inclusion of epistatic effects in whole and sub-genome additive models reduced the additive variance, indicating a lack of orthogonality or statistical independence between additive and epistatic effects. Despite the implementation of recent developments for orthogonal estimations i.e. NOIA and correction for LD and covariance between genetic effects, the lack of independence due to non-orthogonal partitioning of genetic effects could be attributed to the strong linkage disequilibrium, which introduces non-independence between loci (Vitezica et al. 2017). Under the presence of linkage disequilibrium, covariance between genetic effects can be introduced and partitioning genetic effects is not possible (Hill and Mäki-Tanila 2015; Zeng et al. 2005). In simulated and experimental studies, lack of orthogonality was evidenced through a high correlation between additive and epistatic variance component estimates (Vitezica et al., 2017; Raffo et al., 2022). Lack of orthogonality has also been observed in other studies within the context of hybrid breeding (González-Diéguez et al. 2021; Bernardo 1995). González-Diéguez et al. (2021) reported that additive genomic relationship tends to capture additive-by -additive effects when there is a significant correlation between additive and additive-by-additive genomic relationships. This finding aligns with the explanation given by Raffo et al. (2022) and the observations from our study, which reported a correlation of 0.55 between G and I. Furthermore, epistasis itself also contributes to additive genetic variance (Hill et al. 2008; Mäki-Tanila and Hill 2014; Vitezica et al. 2017). Although we did not obtain empirically orthogonal estimates, using methods from Sorensen, Fernando, and Gianola (2001) and Lehermeier et al. (2017), we derived genomic variance estimates that, when combined with error variance estimates, match the sample variance of phenotypes. Therefore, even if the partitioning into additive and epistatic genetic effects is biased due to strong linkage disequilibrium (LD), the total genetic variance estimates remain accurate. This checkpoint—summing the estimated genomic variance and error variance—is often overlooked but is crucial to ensure the correctness of genetic variance component estimates.

In this study, the inclusion of epistasis effects in additive models increased the total genetic variance and reduced the deviance information criteria. In contrast, a study of the Nordic Seed A/S program reported a slight reduction in total genetic variance when epistasis was modeled (Raffo et al., 2022), while a study of Cornell’s and CIMMYT’s breeding populations found that partitioning whole genome effects into sub-genome additive and epistatic effects only marginally increased the AIC for wheat grain yield (Santantonio et al., 2019). One possible explanation for differing results could relate to the large genetic diversity of the K-State wheat breeding program. The introduction and maintenance of diverse germplasm in the K-State program is expected to increase the allele frequency of genomic regions that would otherwise be (nearly) fixed in programs with limited genetic diversity, as may be the case for CIMMYT, Cornell and Nordic Seed A/S programs. A small allele frequency can ultimately convert epistatic variance into additive variance (Whitlock et al., 1995; Cheverud and Routman, 1996; Holland, 2001; Technow et al., 2019), while the segregation of epistatic interactions found at a relatively high minor allele frequencies can increase the statistical power to capture epistatic interactions.

### Sub-Genome Additive Effects as a Tool for Parental Selection if Epistasis is Irrelevant

Estimation of sub-genome additive effects has the potential of attributing biological importance to genetic effects commonly assigned to the whole genome (Santantonio et al 2019) and could be leveraged for selecting parental lines with complementary sub-genome effects when designing breeding crosses. A study of Cornell’s and CIMMYT’s wheat breeding populations found that genotypes with the highest sub-genome additive effects were normally not among the top genotypes having high whole genome effects, and a low correlation between sub-genomes was regarded as a potential indicator of independence between sub-genomes (Santantonio et al., 2019). In this study, making breeding crosses using parental lines (genotypes) having highest additive effects within each sub-genome could produce theoretical maximum whole genome effects of 13.7, 21% higher than the observed highest whole genome effect (11.3). The success of this approach is tightly linked to the interplay of epistatic effects between and within sub-genomes. Although low pairwise correlations amongst sub-genomes were observed, high epistatic variance resulted from both intra-and inter-sub-genome interactions, variances which were nearly zero in Cornell’s and CIMMYT’s breeding populations (Santantonio et al., 2019). In addition, estimation of sub-genome additive effects was less accurate than sub-genome additive plus epistatic effects in predicting which lines reached the elite stages of evaluation in the wheat breeding pipeline. Although the magnitude of epistasis is hard to estimate, epistasis has been reported to affect several traits in wheat, including stem rust (Singh et al., 2013; Rouse et al., 2014), stripe rust (Vazques et al., 2015), plant height (Zhang et al., 2008; Sannemann et al., 2018), photoperiod response (Shaw et al., 2020), vernalization (Kippes et al., 2014), and grain yield heterosis (Boeven et al., 2020; Jiang et al., 2017). These reports suggest epistatic effects should not be overlooked when estimating or predicting the genetic value of wheat genotypes.

In the germplasm used in this study, all sub-genomes exhibited similar additive effects on grain yield in wheat, despite differences in the number of markers within each sub-genome. While the D sub-genome is typically associated with limited genetic variability and fewer markers, one might expect a lower contribution from it. However, the results indicate that the D sub-genome contributed similarly to sub-genomes A and B. Bernardo (2020) also observed equal contributions from each sub-genome for grain yield, although contributions varied for other traits, suggesting that the impact may depend largely on the specific trait being analyzed. The high additive effects could be in part attributed to major QTLs conferring resistance to barley yellow dwarf virus (BYD). In 2017, a severe BYD infection affected the experimental farm, but a large proportion of the advanced genotypes had parental lines carrying *Bdv1* or *Bdv2* resistance genes. *Bdv1* reportedly confers partial tolerance to BYD and is located on chromosome 7D (Singh et al. 1993; Ayala et al. 2002), while *Bdv2*, an introgression from *Thinopyrum intermedium*, is also found on chromosome 7D in hexaploid wheat (Banks et al.,1995; Hohmann et al., 1996; Larkin et al., 2002). Hence, the presence of *Bdv1* and *Bdv2* alleles in many advanced families could have drastically increased the genetic effects associated with the D sub-genome, as the high disease pressure greatly affected grain yield performance.

### Prediction Accuracy Improves When Epistasis is Modeled

As epistasis is potentially an important component of genetic effects impacting grain yield, inclusion of epistatic effects in genomic prediction models could significantly improve prediction accuracies. In this study, modeling epistasis in whole or sub-genome models marginally but significantly improved prediction accuracies, contrasting with the results from Cornell’s and CIMMYT’s wheat breeding populations, where no improvement in prediction accuracy was observed for grain yield, even though inclusion of epistatic effects improved predictability of other traits (Santantonio et al., 2019). In a study of CIMMYT’s Global Wheat and Semiarid wheat breeding populations, an average improvement of 6% in prediction accuracy was observed when modeling additive-by-additive epistatic interactions (Jiang and Reif, 2015), while an increase of 5% in prediction accuracy was achieved in KWS European wheat breeding program (He et al., 2016). These consistent but small improvement in prediction accuracy reported do not support the expectation that epistasis has a major role in wheat given the allopolyploid nature of wheat. In allopolyploid species, sub-functionalization could create positive or negative epistasis (Lynch and Force, 2000), competition for the same substrate (Qian et al., 2010), or affect the patterns of gene expression (Leach et al., 2014; Akhunova et al., 2010), and directly influence the expression of agronomic traits. Furthermore, homeologous interaction effects could be minor relative to the total epistatic genetic effect in wheat, since epistasis has been reported in diploid wheat, *Triticum monccum* (Tranquilli and Dubcovsky, 2000), and also within sub-genomes in hexaploid wheat (Reif et al., 2011; Sehgal et al., 2020). One potential reason for the marginal improvement in prediction accuracy when modeling epistasis lies in the highly quantitative genetic architecture of grain yield. Every stage of crop development is influenced by environmental stresses, and all of which add to total expression of grain yield at the end of the breeding cycle. As various genetic interactions are potentially triggered during crop development, the contribution of each component to grain yield may be so small that capturing epistatic effects is extremely difficult.

## Conclusion

To delineate the genetic architecture of wheat, as expressed for grain yield, whole genome additive and epistatic effects were estimated at the whole genome level and at the sub-genome level. We implemented recent developments and methods for estimating genetic variance components including the natural and orthogonal interaction (NOIA) approach to account for deviation from HWE and correction for LD and covariance between genetic effects. Although estimated phenotypic variance matched the observed phenotypic variance, partitioning of genetic variance components into additive and epistatic variance components was not orthogonal, which can be attributed to the linkage disequilibrium. Incorporating epistasis in additive models did augment the total genetic variance and reduced the DIC, indicating existence of epistatic effects controlling the expression of the trait grain yield. Using a fully additive model, additive effects were estimated for each sub-genome with results indicating that genotypes with largest whole genome additive effects also tended towards intermediate to large sub-genome additive effects. Estimating sub-genome additive effects has potential as a tool during parental selection, as it could permit pairing lines with complementary sub-genome effects. However, the success of this approach is tightly linked to the extent of inter and intra sub-genome epistatic effects. Modeling epistasis in genomic selection models marginally but significantly improved prediction accuracy for both whole-genome and sub-genome models, which can potentially contribute to significant improvements in genetic gain over multiple cycles of selection and advancement.

## Data Availability

Supplemental files are available at the Figshare portal. The files Sub-Genome-A.hmp.txt, Sub-Genome-B.hmp.txt and Sub-Genome-D.hmp.txt refer to the molecular marker data partitioned by each sub-genome of hexaploid wheat. The BLUPs.txt file refers to the grain yield BLUPs. The Var.R, GS.R and Rank.R are the R codes used to estimate the variance components, to run the genomic prediction models, and to rank lines according to sub-genome additive effects, respectively.

