## Supplemental Tables for "Improving genomic selection in hexaploid wheat with sub-genome additive and epistatic models"

**Table 1.** Posterior means (± posterior standard deviation) of additive variance ($V_G$), epistatic variance ($V_I$), total genetic variance ($V_T$), residual variance ($V_E$), estimated phenotypic variance ($V_P$), genomic broad-sense heritability ($H^2$) and deviance information criteria (DIC) estimated with models WG-A (whole genome additive) and WG-AE (whole genome additive plus epistatic).

| Model | $V_G$ | $V_I$ | $V_T$ | $V_E$ | $V_P$ | $H^2$ | DIC |
|---|---|---|---|---|---|---|---|
| 1 | 10.21 ± 0.74 | - | 10.21 ± 0.74 | 27.99 ± 0.85 | 38.19 ± 0.85 | 0.27 ± 0.02 | 23592.16 |
| 2 | 4.21 ± 0.83 | 11.09 ± 1.19 | 14.96 ± 0.9 | 23.29 ± 0.89 | 38.26 ± 0.81 | 0.39 ± 0.02 | 23346.74 |

**Table 2.** Variance (diagonal), covariance (below diagonal) and correlation (above diagonal) of additive and epistatic effects from whole genome additive and epistatic Model 2.

| | Additive | Epistatic |
|---|---|---|
| Additive | 4.21 | 0.00 |
| Epistatic | -0.17 | 11.09 |

**Table 3.** Posterior means (± posterior standard deviation) of additive variance in sub-genome A ($V_A$), B ($V_B$) and D ($V_D$), intra genomic epistatic variance in sub-genomes A ($V_{AA}$), B ($V_{BB}$) and D ($V_{DD}$), inter genomic epistatic variance in sub-genomes AB ($V_{AB}$), AD ($V_{AD}$) and BD ($V_{BD}$), total genetic variance ($V_{TG}$), residual variance ($V_E$), estimated phenotypic variance ($V_P$), genomic broad-sense heritability ($H^2$) and deviance information criteria (DIC) estimated with models SG-A (sub-genome additive) and SG-AE (sub-genome additive plus epistatic).

| Model | $V_A$ | $V_B$ | $V_D$ | $V_{AA}$ | $V_{BB}$ | $V_{DD}$ | $V_{AB}$ | $V_{AD}$ | $V_{BD}$ | $V_G$ | $V_E$ | $V_P$ | $H^2$ | DIC |
|---|---|---|---|---|---|---|---|---|---|---|---|---|---|---|
| 3 | 4.05 ± 0.75 | 3.87 ± 0.71 | 3.94 ± 0.72 | - | - | - | - | - | - | 10.88 ± 0.72 | 27.45 ± 0.8 | 38.33 ± 0.84 | 0.28 ± 0.02 | 23569.99 |
| 4 | 1.42 ± 0.47 | 1.54 ± 0.44 | 1.62 ± 0.47 | 1.76 ± 0.57 | 2.29 ± 0.81 | 2.05 ± 0.74 | 1.67 ± 0.56 | 2.58 ± 0.97 | 2.39 ± 0.99 | 16.33 ± 0.86 | 22.22 ± 0.85 | 38.55 ± 0.8 | 0.42 ± 0.02 | 23289.3 |

**Table 4.** Correlation between whole genome (Model 1) and sub-genome additive effects. The above diagonal shows the correlation of sub-genome additive effects from Model 3 (purely additive) and below the diagonal from Model 4 (additive plus epistatic).

| | A | B | D | Whole |
|---|---|---|---|---|
| A | - | 0.18 | 0.27 | 0.72 |
| B | 0.14 | - | 0.14 | 0.69 |
| D | 0.29 | 0.10 | - | 0.61 |
| Whole | 0.63 | 0.63 | 0.54 | - |

**Table 5.** Mean (standard deviation), maximum and minimum additive effects associated with sub-genome A, B and D (Model 3) and the whole genome (Model 1).

| | A | B | D | Whole |
|---|---|---|---|---|
| Mean | 0 ± 1.27 | 0 ± 1.27 | 0 ± 1.25 | 0 ± 2.51 |
| Maximum | 4.47 | 4.81 | 4.43 | 11.34 |
| Minimum | -4.33 | -3.88 | -4.64 | -10.19 |

**Table 6.** Variance (diagonal), covariance (below diagonal) and correlation (above diagonal) between additive and epistatic effects from sub-genome additive and epistatic Model 4.

| | $V_A$ | $V_B$ | $V_D$ | $V_{AA}$ | $V_{BB}$ | $V_{DD}$ | $V_{AB}$ | $V_{AD}$ | $V_{BD}$ |
|---|---|---|---|---|---|---|---|---|---|
| $V_A$ | 1.42 | 0 | 0 | 0 | 0 | 0 | 0 | 0 | 0 |
| $V_B$ | -0.05 | 1.54 | 0 | 0 | 0 | 0 | 0 | 0 | 0 |
| $V_D$ | 0.02 | -0.06 | 1.62 | 0 | 0 | 0 | 0 | 0 | 0 |
| $V_{AA}$ | -0.04 | -0.01 | -0.01 | 1.76 | 0 | 0 | 0 | 0 | 0 |
| $V_{BB}$ | -0.01 | -0.02 | -0.01 | -0.01 | 2.29 | 0 | 0 | 0 | 0 |
| $V_{DD}$ | -0.01 | -0.01 | -0.03 | -0.01 | -0.01 | 2.05 | 0 | 0 | 0 |
| $V_{AB}$ | 0.00 | 0.00 | -0.01 | -0.02 | -0.01 | -0.01 | 1.67 | 0 | 0 |
| $V_{AD}$ | -0.01 | -0.01 | -0.02 | -0.02 | -0.01 | -0.01 | -0.01 | 2.58 | 0 |
| $V_{BD}$ | -0.01 | -0.01 | -0.02 | -0.01 | -0.02 | -0.01 | -0.01 | 0.00 | 2.39 |

**Table 7.** Correlation between total genetic effects (that is the sum of additive and epistatic effects, if epistasis was modeled) of Models 1-4.

| | **Model 1** | **Model 2** | **Model 3** | **Model 4** |
|---|---|---|---|---|
| Model 1 | - | 0.92 | 0.98 | 0.90 |
| Model 2 | | - | 0.91 | 0.99 |
| Model 3 | | | - | 0.91 |
| Model 4 | | | | - |

**Table 8.** Correlation between additive effects of Models 1-4.

| | **Model 1** | **Model 2** | **Model 3** | **Model 4** |
|---|---|---|---|---|

| | | | | |
|---|---|---|---|---|
| Model 1 | - | 0.91 | 0.99 | 0.89 |
| Model 2 | | - | 0.89 | 0.97 |
| Model 3 | | | - | 0.90 |
| Model 4 | | | | - |
